# Modeling neuronal avalanches and long-range temporal correlations at the emergence of collective oscillations: continuously varying exponents mimic M/EEG results

**DOI:** 10.1101/423921

**Authors:** Leonardo Dalla Porta, Mauro Copelli

**Affiliations:** Departamento de Física, Universidade Federal de Pernambuco (UFPE), Recife, PE, Brazil.

## Abstract

We revisit the CROS (“CRitical OScillations”) model which was recently proposed as an attempt to reproduce both scale-invariant neuronal avalanches and long-range time correlations. With excitatory and inhibitory stochastic neurons locally connected in a two-dimensional disordered network, the model exhibits a transition from an active to an oscillating state. Precisely at the transition, the fluctuations of the network activity have detrended fluctuation analysis (DFA) exponents close to one, and avalanches (defined as supra-threshold activity) have power law distributions of size and duration. By simulating larger system sizes, we show that, differently from previous results, the exponents governing the distributions of avalanche size and duration are not necessarily those of the mean-field directed percolation universality class (3/2 and 2, respectively). Instead, exponents obtained via a maximum-likelihood estimator vary continuously in a narrow region of parameter space. Around that critical region, moreover, the values of avalanche and DFA exponents display a spread with negative correlations, in qualitative agreement with the interindividual variability that was experimentally observed in M/EEG data.

## 1 Introduction

The critical brain hypothesis has emerged in the last decades as a potential framework for theoretically addressing many intriguing questions that have challenged neuroscientists. In light of the nonlinear nature of individual neuronal dynamics and our rudimentary understanding of the collective phenomena that emerge when they interact, these questions include, for instance, the issue of segregation versus integration, optimization of dynamical repertoire, and response to external stimuli, among others (see e.g. Refs. [4, 28] for reviews).

A seminal work by Beggs and Plenz in 2003 reported neuronal avalanches experimentally recorded in vitro [2], lending support to the criticality hypothesis and effectively laying the groundwork for a vast field of research that has involved neuroscientists and physicists alike [23]. In their original setup, spontaneous local field potentials were recorded from cultured slices of the rat brain [2]. The term “neuronal avalanche” seemed like a natural choice, since the observed bursts of activity were interspersed by silence and showed a clear separation of time scales (their duration being much shorter than the interavalanche intervals). Moreover, their size s, defined as either the number of electrodes with suprathreshold activity or the sum of the potentials, were shown to be statistically distributed according to a power law, *P*(*s*) ~ *s*^−1.5^ [2]. Such scale-invariant statistics are one of the hallmarks of a critical system [32, 18].

Other signatures of criticality in the brain have been proposed, such as long-range time correlations, which have been observed both at macroscopic and microscopic levels. For instance, detrended fluctuation analysis (DFA) [21] performed in electroencephalographic and magnetoencephalographic signals have revealed temporal correlations and power-law scaling behaviour in spontaneous oscillations of the normal human brain during large time scales [17, 12]. At a much smaller scale, a similar technique showed that spike avalanches recorded in different regions of freely-behaving rats also have long-range time correlations [25].

A natural next step from the modeling perspective would be to conjugate both ideas. The search for a model that can produce both power-law distributed avalanches *and* long-range time correlations, however, is likely to face some theoretical challenges. Let us briefly discuss why.

It is important to remember that one of the appeals of the 3/2 exponent experimentally observed by Beggs and Plenz is that it coincides with the critical exponent for a branching process [13], which has therefore become a theoretical workhorse in the field. Or, if one extends the idea to rather general networks [16], it coincides with the critical exponent of any model belonging to the universality class of directed percolation (DP) in dimension larger than or equal to its upper critical dimension (*d_c_* = 4). In other words, *τ* = 3/2 is a mean-field exponent for avalanche-size distributions of a class of models in which a continuous phase transition occurs between an absorbing and an active phase [18, 19].

A minimum model of this universality class consists of a large (ideally infinite) number of units (“neurons” or “regions of interest”) which can be either “on” (active) or “off” (inactive) [18]. If a unit is “off”, it may switch to an “on” state with a probability that increases with the number of “on” neighbors and some coupling, say, *λ*. If a unit is “on”, it spontaneously switches to “off” at some constant rate (which accounts for the typical duration of a spike, for instance; variants may include additional states to model the refractory period [14]). The absorbing phase bears its name because, for sufficiently small *λ*, eventually all units go to the “off” state and the dynamics of the whole network stops. For sufficiently large *λ*, on the other hand, propagation of the “on” state among units occurs at such a rate that the system stays in an active phase, characterized by stable self-sustained activity, i.e. nonzero time- and ensemble-average density of active sites 〈*ρ*〈 (the typical order parameter for these models). The boundary in parameter space between those two qualitatively different regimes is the critical point *λ* = *λ_c_*, above which 〈*ρ*〉 departs continuously from zero as 〈*ρ*〉) ~ (*λ* – *λ_c_*)^*β*^, where *β* is a critical exponent [18]. Precisely at *λ* = *λ_c_*, the system is on the verge of displaying self-sustained activity, so perturbations to the absorbing state (which is still stable) do not have a characteristic time to die out or a characteristic size. In fact, their size *s* (defined as the number of active sites along the excursion) and duration *d* (defined as the time between the first and last active site in between epochs of complete quiescence) are power-law distributed at the critical point: *P*(*s*) ~ *s*^−*τ*^ and *P*(*d*) ~ *d*^−*τ_d_*^. These critical exponents depend on network dimensionality *D* [19], and for *D* ≥ 4 the mean-field exponents are precisely *τ* = 3/2 and *τ_d_* = 2, as observed by Beggs and Plenz [2].

In the model, these perturbations are called “avalanches”, among other reasons, because their dynamics are subject to an infinite separation of time scales *by construction*: once an avalanche is over (all sites “off”), the next one will not start unless the system is arbitrarily perturbed again. This unassuming detail, however, poses a challenge if one wants to explore this class of models to reproduce the above mentioned long-range time correlations which were observed experimentally [17, 25]. Since consecutive avalanches are, by definition, separated by returns to the absorbing state, it is not apparent how inter-avalanche correlations could emerge (self-organizing mechanisms are a potential candidate, yet to be tested [15]).

Given this state of affairs, an interesting perspective was put forward by the CROS (“CRitical OScillations”) model proposed by Linkenkaer-Hansen *et al* [24, 11]. The model has both excitatory and inhibitory stochastic spiking neurons arranged in an *L* × *L* square lattice, each neuron interacting with a controlled random fraction of its neighbors within a small *ℓ* × *ℓ* region (Fig. 1A, see Methods for details). All excitatory neurons receive an independent constant Poisson input which, contrary to the DP universality class we discussed previously, ensures the model does *not* have an absorbing state. Changing the model parameters controlling excitation and inhibition (*r*_E_ and *r*_I_), Linkenkaer-Hansen *et al* reported instead a transition from a regime of constant activity to one of collective oscillations, as marked by the emergence of a peak in the Fourier spectrum of the global network activity. At the transition line in the (*r*_E_,*r*_I_) plane, long-range time correlations were observed with DFA exponents *α* ⋍ 1, in contrast with uncorrelated fluctuations far from the transition (*α* ⋍ 0.5). Remarkably, at the *same* transition line Linkenkaer-Hansen *et al* also managed to obtain avalanches with power law distributions of size [24] and duration [11], in either case with the respective mean-field exponent *τ* = 3/2 and *τ_d_* = 2. Importantly, in the absence of an absorbing state, their definition of an avalanche required the imposition of an *ad hoc* threshold *θ*, defined as a fraction Γ of the median network activity *m̃*. In the CROS model, therefore, it is the crossing of an arbitrary threshold upwards and downwards that marks respectively the beginning and end of an avalanche (see Fig. 2A and Methods). Differently from a DP-like model with an absorbing state, the system dynamics itself controls when the threshold will be crossed. And that is the ingredient allowing for long-range time correlations.

**Figure 1:**
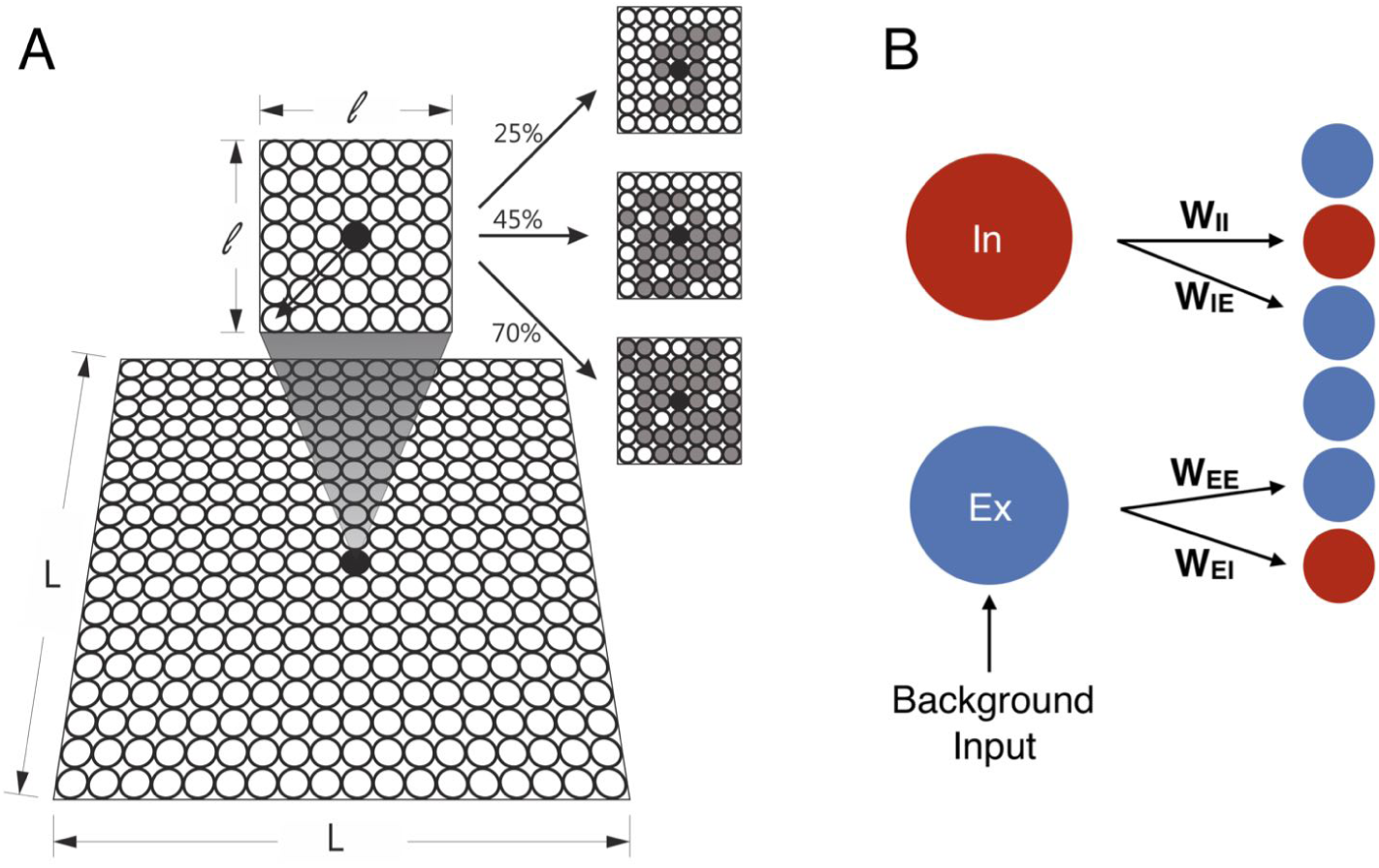
CROS model. (A) The model consists of excitatory (Ex) and inhibitory (In) neurons arranged in a *L* × *L* square lattice with open boundaries. Each neuron may connect locally to a random fraction of its neighbors within a *ℓ* × *ℓ* square (see Methods). In gray we exemplify the neurons connected to a central neuron (black dot) with connectivity of 25% (top), 45% (middle), and 70% (bottom). (B) The connection weights (*W_ij_*) are fixed and depend only on the nature of the presynaptic (*i*) and postsynaptic (*j*) neurons.

**Figure 2:**
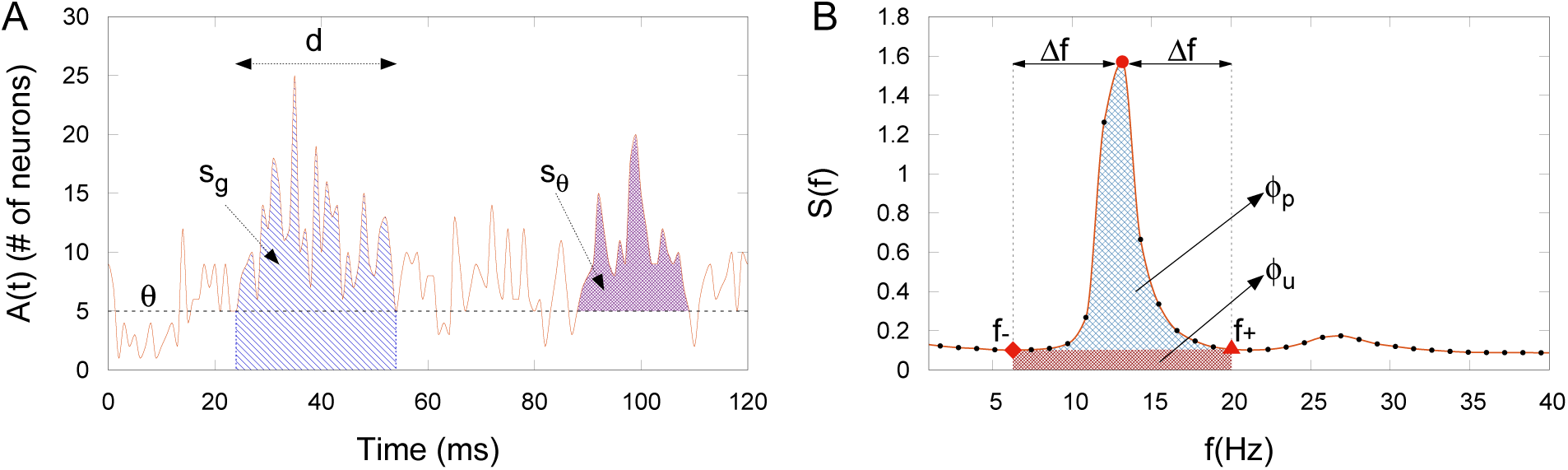
Neuronal avalanches and order parameter definitions. (A) A neuronal avalanche starts and ends when the fluctuations of the integrated network activity cross a threshold *θ*. The size of an avalanche can be defined as s_*g*_, the total number of spikes, or s_*θ*_, the number of spikes minus the threshold value. The avalanche’s duration *d* is the time that the fluctuations stay above *θ*. (B) Detail of the power spectrum around the region where a peak appears. The order parameter *φ* is given by the ratio between the power peak area *ϕ*_p_ and the total area *ϕ*_p_ + *ϕ*_u_. Red symbols represent the frequencies that bound the *φ* area: *f*_min_(diamond), *f*_peak_(circle) and *f*_max_ (triangle).

The results of the model proposed by Linkenkaer-Hansen *et al* are remarkable in that they seem to reconcile the emergence of long-range time correlations with avalanche distributions which in principle appear in models of a very different nature, as discussed above. The reconciliation is even more appealing given that the coexistence of these two phenomena has been reported experimentally in M/EEG recordings [20, 31]. However, from the theoretical point of view, the model raises several questions, which we set out to investigate here. Firstly, we introduce a tentative order parameter to better characterize the phase transition, while also simulating larger system sizes (*L* = 300) than in the original results (*L* = 50) [24]. Secondly, we point out that it is highly counterintuitive to have a two-dimensional system exhibiting mean-field exponents. That could be due to an insufficiently small ratio *ℓ*/*L* (which is addressed here, again, by increasing *L*). We argue that, if the onset of oscillations in a two-dimensional network is to be reconciled with DP critical exponents, then the DP exponents for *D* = 2 should also be tested. Thirdly, we explore parameter space in more detail around the transition line, allowing the exponents to be adjusted by a maximum-likelihood estimator (MLE) and testing whether the model satisfies other scaling relations. Finally, we explore the extent to which the model is able to reproduce scaling exponents observed experimentally in M/EEG recordings.

## Methods

### CROS network model

We employed essentially the same CROS (CRitical OScillations) model as proposed by Poil *et al* [24]. Excitatory (80%) and inhibitory (20%) neurons were randomly arranged in a bidimensional *L* × *L* square lattice, with open boundaries (Fig. 1A). Each neuron has a finite local range of connections limited to a square of size *ℓ* × *ℓ* centered around it (Fig. 1A). We used *ℓ* =7 like in the original model [24], which means that a typical neuron (far away from the borders) could connect to a maximum of 48 other neurons in their neighborhood. Excitatory (*r*_E_) and inhibitory (*r*_I_) connectivity are the two free parameters of this model, being set between 2 – 60% and 30 – 90%, respectively, of the total number of neurons within the local range. Each neuron sends its outgoing synapses to other neurons at a distance *r* with probability *P*(*r*) = *Ce*^−*r*/*a*_0_^, where *a*_0_ = 1 is the nearest-neighbor distance, and *C* a normalization constant such that ∑_neighbors_ *P*(*r*) equals *r*_E_ or *r*_I_ for presynaptic excitatory and inhibitory neurons, respectively. Notice that the networks have disorder, in the sense that, for a given point (*r_E_*,*r_I_*) in parameter space, different synaptic connections are possible (corresponding to how the specific grey neurons in Fig. 1A are randomly selected within the *ℓ* × *ℓ* interaction range). We will return to this point in the discussion of the results. Five networks for each combination of *r_E_* and *r*_I_ were created and simulated for 2^20^ time steps (*dt*) of 1 ms. Each connection had a weight (*W_ij_*) depending on the nature (excitatory or inhibitory) of the presynaptic and postsynaptic neurons (Fig. 1B).

The neuron dynamics starts with a neuron *i* integrating all the received input from the connected neighbors and updating its synaptic current *I_i_*, which is also subject to an exponential synaptic decay:

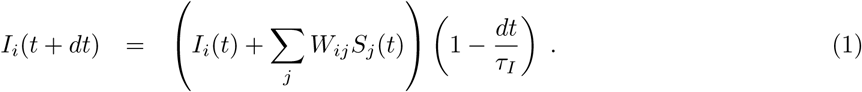

In Eq. (1), *S*(*t*) ∈ {0,1}^N^ is a binary vector denoting whether the presynaptic neurons fired in the previous time step, whereas the connection weights are fixed at *W_1E_* = 0.011, *W_EE_* = 0.02 and *W_E1_* = *W_II_* = –2 and the time constant is *τ_I_* = 9 ms [24]. Given the synaptic current *I_i_* we compute *R_Si_*, which is also subject to an exponential decay:

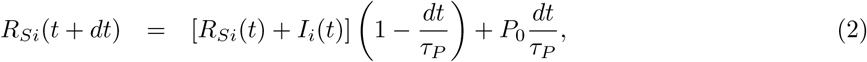

where *τ_P_*(excitatory)= 9 ms, *τ_P_*(inhibitory)= 12 ms and *P*_0_ is the background spiking probability with *P_0_*(excitatory) = 10^−6^ and *P*_0_(inhibitory) = 0 [24]. Each neuron spikes with probability

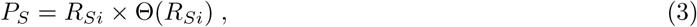

where Θ is the Heaviside function. Then *S* is updated for the next time step. If a neuron spikes we reset *R_S_* to –2 (excitatory neurons) or –20 (inhibitory neurons) [24].

### Neuronal avalanches

Given that some balance of *r_E_*:*r*_I_ produce networks that are continuously active, a threshold is necessary to define the start and end of an avalanche. Poil et. al. [24] proposed a threshold defined as 50% of the median spike activity of the network. Here we defined the threshold *θ* as:

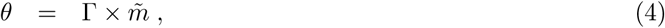

where Γ ∈ [0,1] is a number and *m̃* the median activity (so the original CROS model used Γ = 0.5). A neuronal avalanche starts and ends when the fluctuation of the summed activity of the network *A*(*t*) = ∑_*i*_ *S_i_*(*t*) crosses *θ*. If an avalanche starts at time *t_i_* and ends at time *t_f_*, its duration is *d* = *t_f_* – *t_i_* (Fig. 2A). We studied two definitions for the size of an avalanche: i) 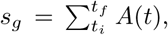 i.e. the total area under the *A*(*t*) curve, as originally proposed by Poil et *al.* [24]; and ii) 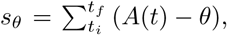 i.e. the area above the threshold, as recently proposed by Del Papa et *al.* [7].

### *κ* index

We used the *κ* index proposed by Shew *et al* [29] to obtain a first approximate assessment of how close an observed (simulated) distribution *P^obs^* is from a known theoretical distribution *P^th^*. We focused on power laws distributions of avalanche size and duration (both referred to simply as *x* in what follows). If we restrict the analysis of a power law distribution *P^th^*(*x*) ~ *Cx*^−*μ*^ to an interval [*x_min_*, *x_max_*], in the continuum limit one can calculate both the normalization constant 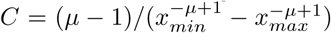 and the theoretical cumulative function 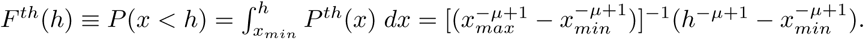

Let 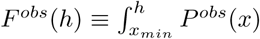 *dx* be the observed cumulative function. The *κ* index, which quantifies the difference between an observed and a theoretical distribution, is defined as

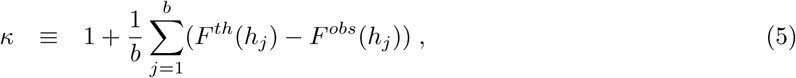

where *h_j_* are *b* = 10 observables logarithmically spaced between *x_min_* and *x_max_*. With this definition, one obtains *κ* ⋍ 1 for an observed distribution which is close to the theoretical power law, *κ* < 1 for a subcritical (e.g. exponential-like) distribution and *κ* > 1 for a supercritical distribution (say, with a characteristic bump at larger values of *x*).

In our case *μ* stands for *τ* and *τ_d_*, critical exponents from the Direct Percolation (DP) universality class for size and duration of an avalanche, respectively. These exponents depend on the dimension of the underlying model. As mentioned previously, *τ* = 3/2 and *τ_d_* = 2 stand for DP models at or above the upper critical dimension (*d* ≥ *d_c_* =4), i.e. are mean-field exponents. Since the CROS model studied here is in principle twodimensional, we also employ *τ* = 1.268 and *τ_d_* = 1.450, which are the critical exponents for two-dimensional DP [19]. We denote the indices to quantify the scale invariance of avalanche size *s_g_* or *s_θ_* and avalanche duration *d* as *κ_g_*, *_θ_* and *κ_d_*, respectively. Each of these three indices come in two variants, one employing mean-field (MF) exponents for the theoretical distributions and another employing their two-dimensional (2D) counterparts. We present heat maps in parameter space of the different *κ* indices, averaged over 5 realizations of the disorder.

### Maximum Likelihood Estimator

Estimating the quality of a power-law fit with the *κ* index has the drawback that the exponent *μ* must be known *a priori.* In order to loosen this constraint (while complying with more firmly grounded criteria for fitting power-law distributions [5]), we employ the maximum likelihood estimator (MLE) as made available in the `powerlaw` python package [1]. The exponents for size and duration of avalanches were estimated from a power law with an exponential cutoff, *P*(*x*) ~ *x*^−*μ*^ exp(–*x*/*x*_0_), with *μ* and *x*_0_ as free parameters and fixing minimum values *s_min_* = 10 and *d_min_* = 4, respectively.

The package supports different probability distributions, which allowed us to use the loglikelihood ratio test [33] to compare the goodness of the fit of the exponentially truncated power-law with that of other possible distributions. We made comparisons with pure power-law, lognormal and exponential distributions. For our data (Figs 4, 5 and 6), the exponentially truncated power-law was always a better fit than those three other distributions, for size as well as for duration of neuronal avalanches. Exceptionally for Fig. 8 we did not test which probability distribution would be the most appropriated and *τ* and *τ_d_* were estimated following an exponentially truncated power-law distribution.

### Scaling relation between avalanche size and duration

At criticality, one expects a power-law relation between the average size 〈*s*〉 of an avalanche of a given duration *d*:

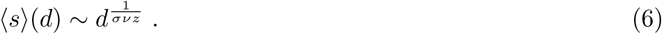

Moreover, the exponents are related to the those ruling the distributions of avalanche size and duration [27, 9, 26]:

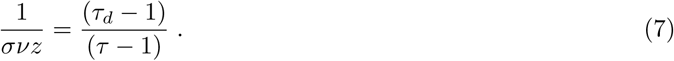

These scaling relations were observed in *in vitro* and *ex vivo* experiments [9, 30]. We probe the model by independently testing whether Eqs. 6 and 7 hold near its critical line.

### Order parameter, power spectrum and DFA analysis

The frequency spectrum was obtained via the Fast-Fourier Transform (FFT), normalized by the network size (*L* × *L*) and the total time of simulation. The spectra were subsequently smoothened by averaging over nonoverlapping windows of 1200 frequency steps (each step with width of 9 × 10^−4^ Hz, resulting in one smoothened point every ~ 1.14 Hz).

The order parameter *φ* was defined as the ratio between the area of the power peak *ϕ*_p_ divided by the total area *ϕ_p_* + *ϕ_u_* (Fig. 2B):

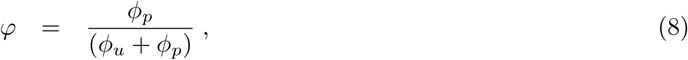

where *ϕ_u_* is the area under the power peak. *φ* is only different from zero if we can detect a peak in power-spectrum. Operationally, once we have the peak frequency *f_p_*, we detected the local minimum *f*_–_ below it (between 4 –18 Hz), which yields Δ*f* ≡ *f_p_* – *f*_–_. Using a symmetrical interval around *f_p_*, we calculated an upper frequency *f*_+_ ≡ *f_p_* + Δ*f*. Given these three points, we fitted a line between *f*_–_ and *f*_+_: the area above the line is *ϕ_p_* and the area under it is *ϕ_u_*.

DFA analysis was performed on the activity time series *A*(*t*) following the standard procedures described in [21, 22, 10]. Briefly, the occurrence of a self-similarity exponent *α* in the time series is a reliable indicator of long-range time correlation if 0.5 < *α* < 1, whereas *α* ~ 0.5 results from uncorrelated fluctuations. If the power spectrum decays as *S*(*f*) ~ 1/*f^v^*, then *v* = 2*α* – 1 [3]. So, when *α* = 1 we have the presence of 1/*f* noise. In this work we estimated *α* from a fixed time range between 4 s and 400 s.

### Experimental M/EEG recordings

To compare the model results with those of experimental M/EEG recordings, the latter data was extracted directly from Ref. [20].

## Results and Discussion

### System-size and threshold dependence

We start by addressing the dependence of the results on the system size *L*. The original results of Poil *et al* were obtained with *L* = 50 and *ℓ* = 7 [24]. As such, the two-dimensional nature of the network is debatable at best.

Note that in the CROS model the definition of an avalanche depends on the threshold parameter *θ* (Fig. 2A), whereas other signatures of criticality (DFA exponents, power spectra and order parameter) do not (Fig. 2B). We therefore initially investigate these two types of markers of criticality separately, starting by those which do not depend on *θ*. To probe the robustness of the results, we kept *ℓ* = 7 while increasing system size up to *L* = 300 (Fig. S1) In all cases, we obtained the same scenario, which we summarize for the largest system size in Fig. 3. Keeping *r*_I_ fixed and increasing *r*_E_, one observes not only an increase in the average activity 〈*A*〉, but eventually also the emergence of oscillations with a characteristic time scale in *A*(*t*) (Fig. 3A). These can be characterized by power spectra, in which a peak emerges (Fig. 3C) at a critical region (line) in the (*r_E_*, *r*_I_) parameter plane (Fig. 3B). Alternatively, the order parameter *φ* can be used to estimate the transition line (Fig. 3D), where its fluctuations are also maximal (Fig. 3E).

**Figure 3:**
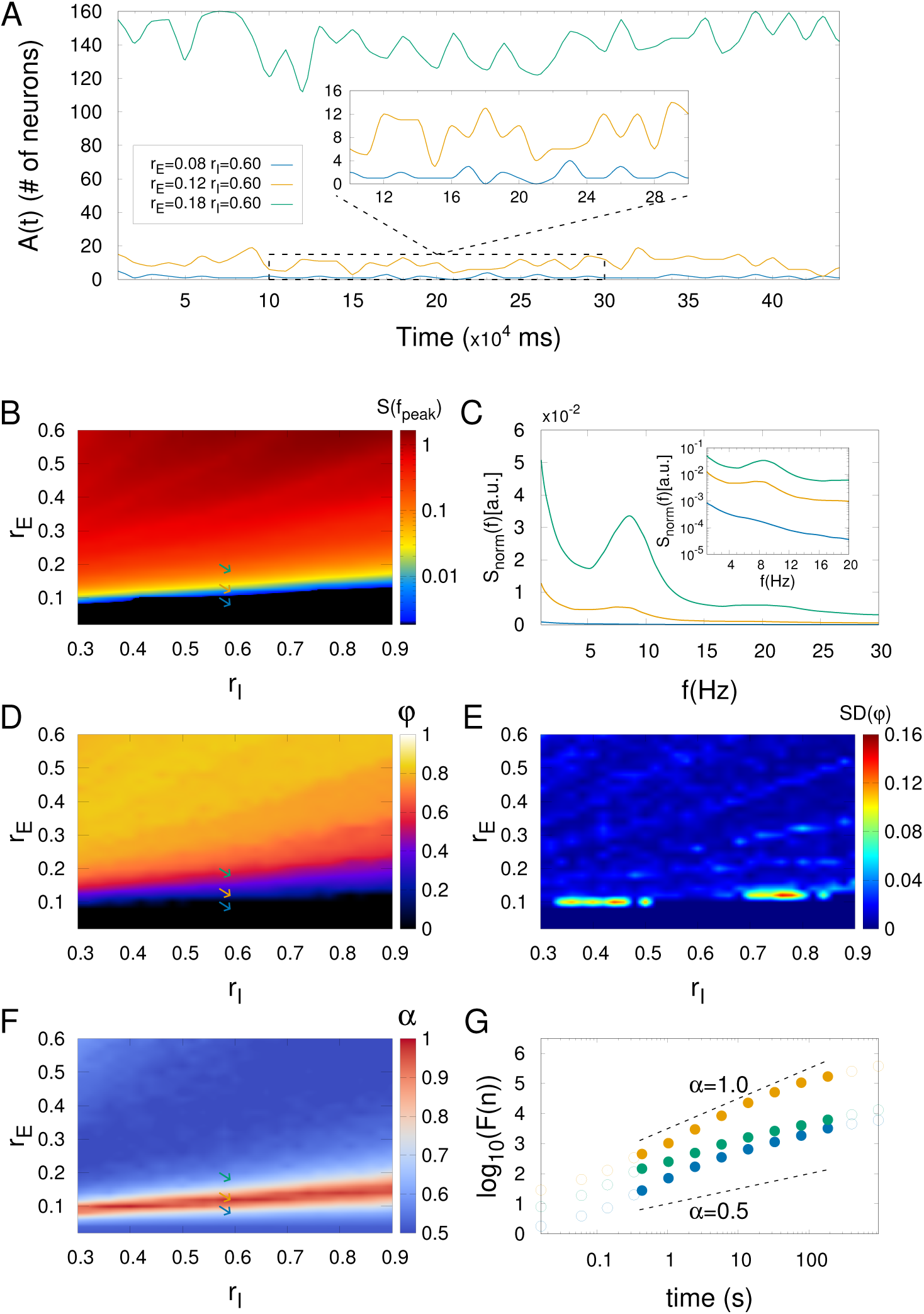
CROS model dynamics with *L* = 300 and *ℓ* = 7. (A) Network activity for three different regimes (balance between excitation and inhibition): low (blue), intermediate (yellow) and high (green). Inset: zoom in the lowest level of activity. (C) Power spectrum of *A*(*t*) for the three regimes described in (A). Inset: zoom in low frequencies with double logarithmic scale. The parameter space suggests a phase transition as shown in heat maps of (B) the power of collective oscillations and (D) the order parameter. Also (E) a larger standard deviation of the order parameter is observed near the critical region. (F) In the same critical region, stronger long-range temporal correlation is indicated by the DFA exponent *α* (see Methods). (G) The DFA exponent *α* is close to one near the critical regime (yellow dots, *α* = 0.97) while subcritical (blue dots, *α* = 0.68) and supercritical (green dots, *α* = 0.62) regimes are close to a white noise (*α* ≈ 0.5). Solid circles indicate the fitting range where *α* was estimated (see Methods). Arrows in (B), (D) and (F) represent parameters described in (A).

For each of these time series, we have calculated the DFA exponent *α*. Interestingly, the critical region is also characterized by larger values of *α* (Figs 3F and 3G), as originally reported [24]. These threshold-independent markers of criticality therefore seem to be robust to an increase in system size, suggesting a phase transition at which the system has long-range temporal correlations while oscillations start to emerge. In what follows, all results were obtained for networks with *L* = 300.

Turning our attention now to avalanches, we investigate the dependence of the results on the threshold *θ* = Γ*m̃* (see Eq. 4). The *κ* index is a useful tool to summarize how close the size and duration distributions are to power laws in parameter space (*as long as* one is confident about which exponent to expect, as we will discuss below). We used *κ_g_* and *κ_θ_* for the original (*s_g_*) and above-threshold-only (*s_θ_*) definitions of avalanche size, respectively (see Methods), with *κ_d_* characterizing the distributions of avalanche durations (*d*). For each of these three, we imposed either the usual mean-field (*τ* = 3/2, *τ_d_* = 2) or the 2D (*τ* = 1.268, *τ_d_* = 1.450) exponents of the directed percolation universality class, in a total of six variants of *κ*. Exploring their heat map in parameter space for Γ = 0.5, 0.75 and 1 (Fig. S2), we obtain critical regions in parameter space where *κ* ⋍ 1, pointing to power-law distributions of sizes and durations. Since the results for *s_θ_* were more robust than for *s_g_*, the former will be used throughout. Except for the largest value of Γ, the phase diagrams also proved to be rather insensitive to this parameter (Fig. S2), so we will use Γ = 0.5 (as originally proposed [24]) unless otherwise stated.

### Power-law distributed avalanches: exponents from 2D or mean-field directed percolation?

With a larger system size (*L* = 300) and local interactions (*ℓ* = 7), we turn to the question whether the avalanche distributions produced by the model are compatible with the universality class of two-dimensional directed percolation (2D-DP). Starting with size distributions, if one imposes the 2D-DP exponent *τ* = 1.268 and calculates the corresponding *κ_θ_*, a critical region with *κ_θ_* ⋍ 1 is indeed found (Fig. 4A). However, if the mean-field directed percolation (MF-DP) exponent *τ* = 3/2 is used instead, the heat map of the corresponding *κ_θ_* index *also* displays a critical region (Fig. 4B) near the 2D-DP one. In both cases, a close inspection of a few representative points in parameter space (Figs 4C and 4D) shows clear subcritical distributions (consistent with *κ_θ_* < 1) as well as supercritical distributions with a mild bump (*κ_θ_* > 1; moving just a bit further into the supercritical region, one obtains essentially a few giant avalanches - often just one). More importantly, fitting the exponent of a power-law with exponential cutoff (with the MLE method) at the putative critical region, one obtains an exponent reasonably close to the value for 2D-DP at one point in parameter space (Fig. 4C) and another close to that of MF-DP at another (Fig. 4D). Both power laws hold for at least three decades, consistently with the maxima in the respective *κ_θ_* heat maps.

**Figure 4:**
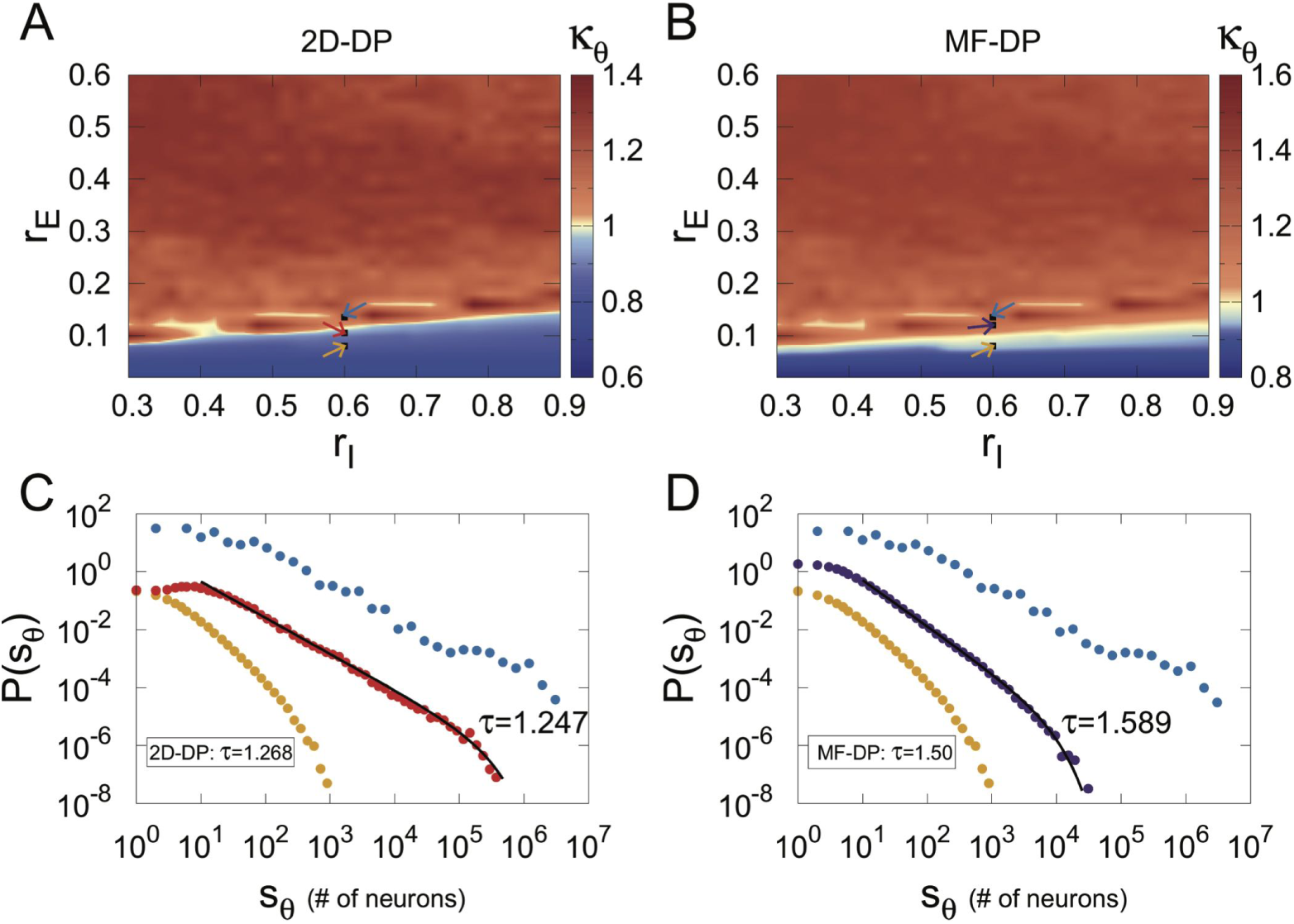
Avalanche size distributions. (A) and (B) show heat maps of the *κ_θ_* index in parameter space employing the *τ*_d_ exponent for the 2D-DP and MF-DP universality classes, respectively. In both cases, avalanches were defined with Γ = 0.5. Arrows indicate the connectivity parameters [*r_E_*:*r*_I_] of networks: [0.080:0.60] (yellow), [0.1225:0.60] (red), [0.1050:0.60] (purple) and [0.1325:0.60] (blue). Representative (single run) distributions for the parameter values indicated by arrows in (A) and (B) are respectively shown in (C) and (D), exemplifying subcritical, critical and supercritical cases.

A very similar scenario is observed when the distributions of avalanche duration *P*(*d*) are studied. Heat maps for the *κ_d_* indices for 2D-DP and MF-DP exponents (Figs 5A and 5B, respectively) *both* show maxima in putative critical regions, with subcritical and supercritical behaviors off these regions. The MLE-fitted exponents are again close to the theoretical 2D-DP and MF-DP values, perhaps with a better fit in the first case in comparison with the second (approximately three decades in Fig. 5C versus two in Fig. 5D).

**Figure 5:**
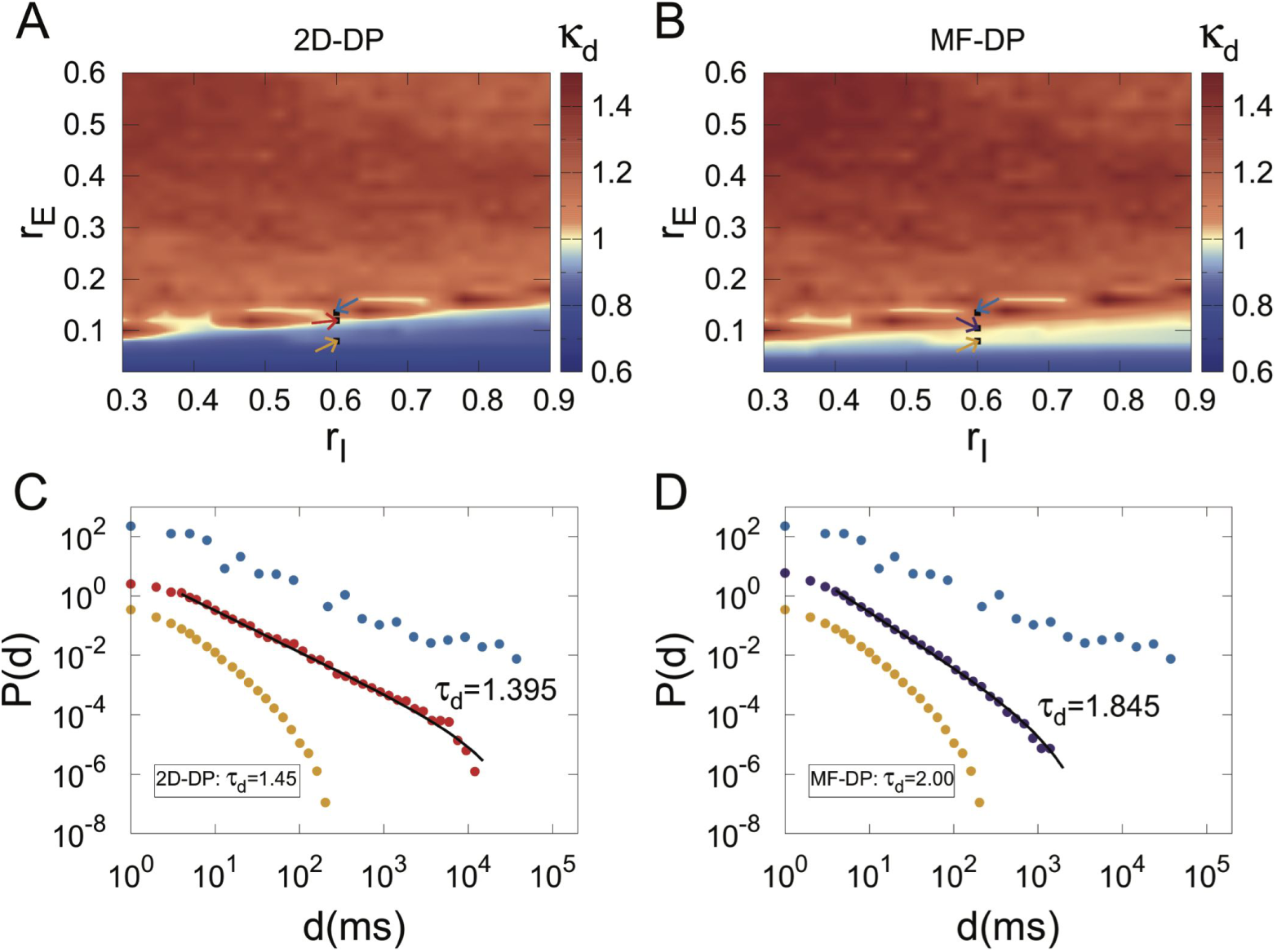
Avalanches duration distributions. (A) and (B) show heat maps of the *κ_d_* index in parameter space employing the *τ_d_* exponent for the 2D-DP and MF-DP universality classes, respectively. In both cases, avalanches were defined with Γ = 0.5. Arrows indicate the connectivity parameters of networks, the same described in Fig. 4. Representative (single run) distributions for the parameter values indicated by arrows in (A) and (B) are respectively shown in (C) and (D), exemplifying subcritical, critical and supercritical cases.

Given the current absence of a proper theoretical connection between a transition to collective oscillations and the DP universality class, the above presented results suggest that the *κ* index is not the most appropriate tool for clarifying this issue, since by construction it relies on *a priori* knowledge of the distribution exponent. Henceforth we therefore relax this constraint, taking a more agnostic approach towards the values of the exponents and letting the MLE method determine them.

### Maximum-likelihood estimator and scaling relations

If one fixes *r*_I_ = 0.6 and measures the order parameter *φ* as the excitatory connectivity *r*_E_ changes, the characteristic plot of a second-order phase transition emerges (Fig. 6A). Around a critical region, *φ* departs continuously from zero and, consistently, the DFA exponent *α* peaks (Fig. 6A corresponds to a cross section of Figs 3D and 3F). In the same region (shaded area in Fig. 6A), the distributions of avalanche sizes are better fitted by power laws with exponential cutoffs than by exponentials or log normals, according to the loglikelihood test (see Methods).

**Figure 6:**
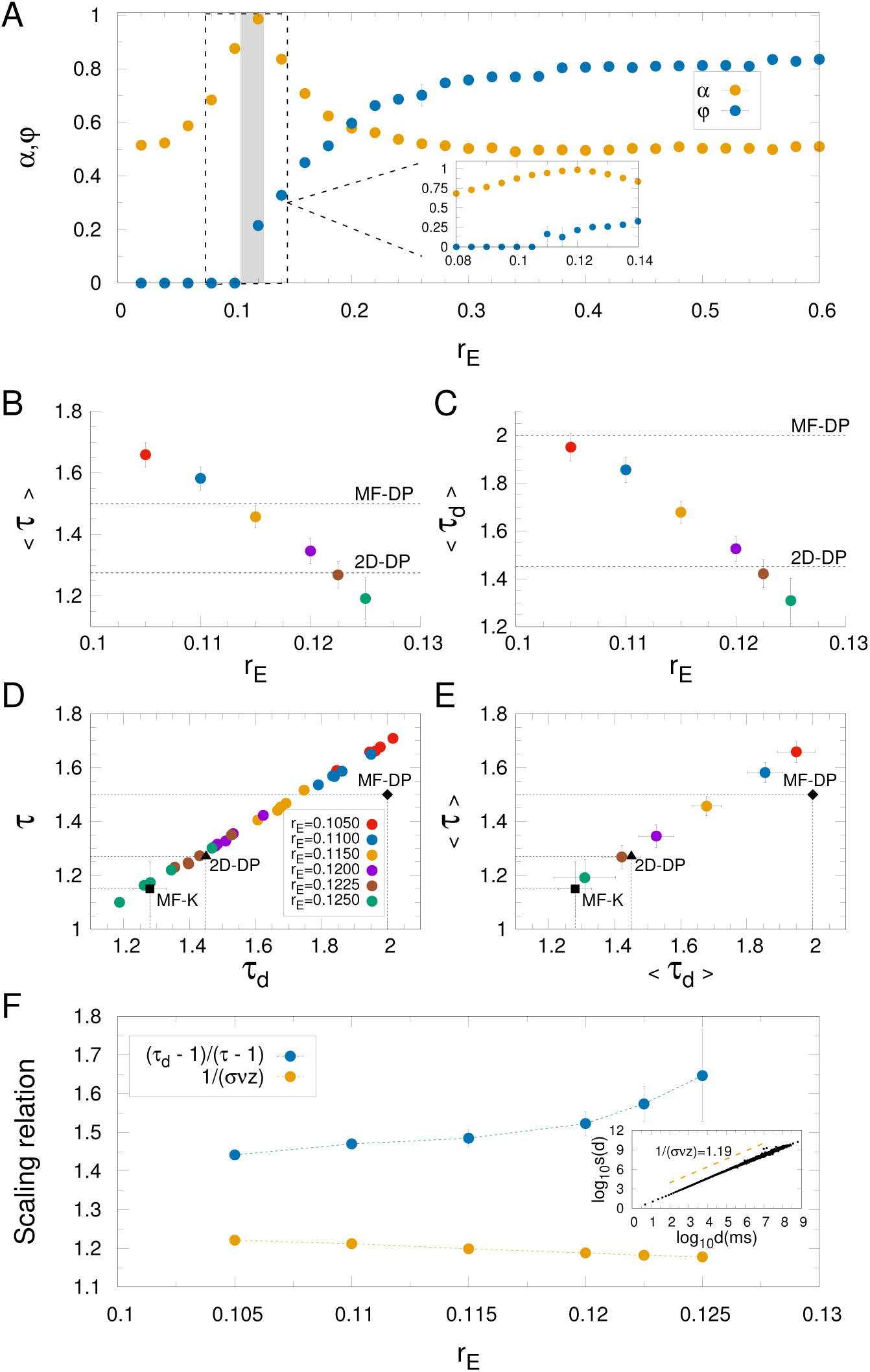
Scaling behavior. (A) DFA exponent (*α*) and order parameter (*φ*) versus *r*_E_, with *r*_I_ = 0.60 fixed. The shaded area represents parameter space where the data was better fitted by a exponentially truncated power-law than exponential or lognormal distributions, according to the log-likelihood ratio test (see Methods). Inset: zoom around the shaded area. (B) and (C): average exponents for avalanche size and duration, respectively. Avalanche size *versus* avalanche duration for (D) 5 different runs and (E) averages over the runs. The black triangle and diamond represent the theoretical exponents for 2D-DP and MF-DP universality classes, respectively, whereas the black square is the result obtained by Coleman *et al* [6] for a variant of the globally coupled Kuramoto model. The same color code applies to (B)-(E). (F) Left-hand (yellow) and right-hand (blue) sides of scaling relation given by Eq. 7 as a function of *r*_E_ in the critical region. Inset: An example of the scaling relation given by Eq. 6 for *r*_E_ = 0.115. In all figures, error bars represent the standard deviation over 5 runs. Avalanches were defined with Γ = 0.5.

Zooming in this region, the MLE method yields exponents for the distributions of size (Fig. 6B) and duration (Fig. 6C) which vary continuously as parameter space is traversed. Note that these are averages over different realizations of the disorder (see Methods). Eventually, both 2D-DP and MF-DP values are crossed for the truncated power laws which best fit the size and duration distributions.

At this point, it is unclear whether either of these two theoretical conjectures should be chosen as a better description of criticality in the model. We therefore applied a more stringent test, asking whether there is some combination of parameters for which the values of *both τ* and *τ_d_* come close to those of either of the two conjectured universality classes. From the functions 〈*τ*〉(*r*_E_) and 〈*τ_d_*〉(*r*_E_) (Figs 6B and 6C), one can parametrically plot 〈*τ*〉 vs 〈*τ_d_*〉, either with (Fig. 6E) or without (Fig. 6D) averaging over the disorder. In the plane (*τ*, *τ_d_*), simulation results eventually come close to the 2D-DP exponents, but not to the MF-DP exponents. This in principle seems consistent with the fact that the model is two-dimensional, especially since care has been taken to increase the system size.

Note also that the precise parameter value at which the two 2D-DP exponents hold depends on the realization of the disorder, with *r*_E_ ranging from 0.1200 to 0.1250 (Fig. 6D). On the one hand, this is not so surprising given that disorder can indeed have strong effects on DP models, including the emergence of a Griffiths region within which dynamical exponents vary continuously [34]. On the other hand, it is unclear to which extent (if any) results for DP models with disorder translate to the CROS model studied here. For instance, Vojta *et al* have simulated two-dimensional contact processes on very large *disordered* lattices, where each bond may or may not exist with a given probability. They have shown that the absorbing-active phase transition is characterized by universal critical exponents [34]. In particular, they have obtained *η* ⋍ 0.15(3) and *δ* ⋍ 1.9(2) [34], which are the exponents governing the growth of the number of active particles (*N*(*t*) ~ *t^η^*) and the survival probability (*P_s_*(*t*) ~ *t*^−*δ*^) in spreading experiments [19]. These exponents are independent of the disorder strength [34]. Rather generic scaling relations have been found for absorbing phase transitions connecting these exponents to those of avalanche distributions, namely [19]: 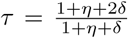 and *τ_d_* = 1 + *δ*. Applying them to the results by Vojta *et al* for disordered two-dimensional systems [34], one obtains *τ* ⋍ 1.62 and *τ_d_* ⋍ 2.9. Although the avalanche size exponent for disordered 2D-DP models lies within the range obtained for the CROS model (Fig. 6B), the avalanche duration exponent clearly does not (Fig. 6C), which seems to exclude a compatibility with the disordered 2D-DP universality class.

If both exponents *τ* and *τ_d_* are simultaneously well fit by the (clean, i.e. without disorder) 2D-DP values at the critical region (Figs 6D and 6E), another consistency relation can be tested. On the one hand, one expects the average avalanche size 〈*s*〉 to scale with the duration *d* as 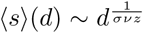 (Eq. 6). We have confirmed that this holds within the critical region of the CROS model (inset of Fig. 6F; as noted previously, however, this relation often holds even far from criticality [9]). On the other hand, there is a scaling relation connecting the exponent in Eq. 6 and those governing the distributions of avalanche size and duration: 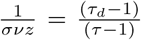 (Eq. 7). Note that the left-hand and right-hand sides of this scaling relation can be independently evaluated, and then compared for consistency. As the parameter *r*_E_ is varied within the critical region (gray strip in Fig. 6A), the left-hand side stays well below the right-hand side, failing to satisfy the scaling relation (Fig. 6F). Note that this failure is independent of a requirement to fit either conjectured universality class (2D-DP or MF-DP): the lines in Fig. 6F do not cross anywhere.

Now we should revisit the dependence of the results on the threshold parameter Γ, a crucial ingredient of the model that directly impacts the very definition of an avalanche (Methods, Eq. 4). While the *κ* index proved to be fairly robust against variations in Γ (Fig. S2), its usefulness was shown to be rather limited. So how does the overall picture of the much more informative Fig. 6 change when the threshold is varied?

First, we observe that the absolute values of both avalanche exponents increase with increasing Γ (Figs. 7A and 7B). This is interesting, because it relates to a similar result originally obtained by Beggs and Plenz [2]: by decreasing the time bin employed to slice their time series, they observed an increase in the absolute value of *τ*. In other words, by imposing stricter conditions (shorter silences), larger avalanches became less likely, in the sense that the power law describing the size distribution became steeper. Our results reproduce the same trend (also seen by di Santo et al. [8]), since imposing larger thresholds is tantamount to decreasing the bin width.

**Figure 7:**
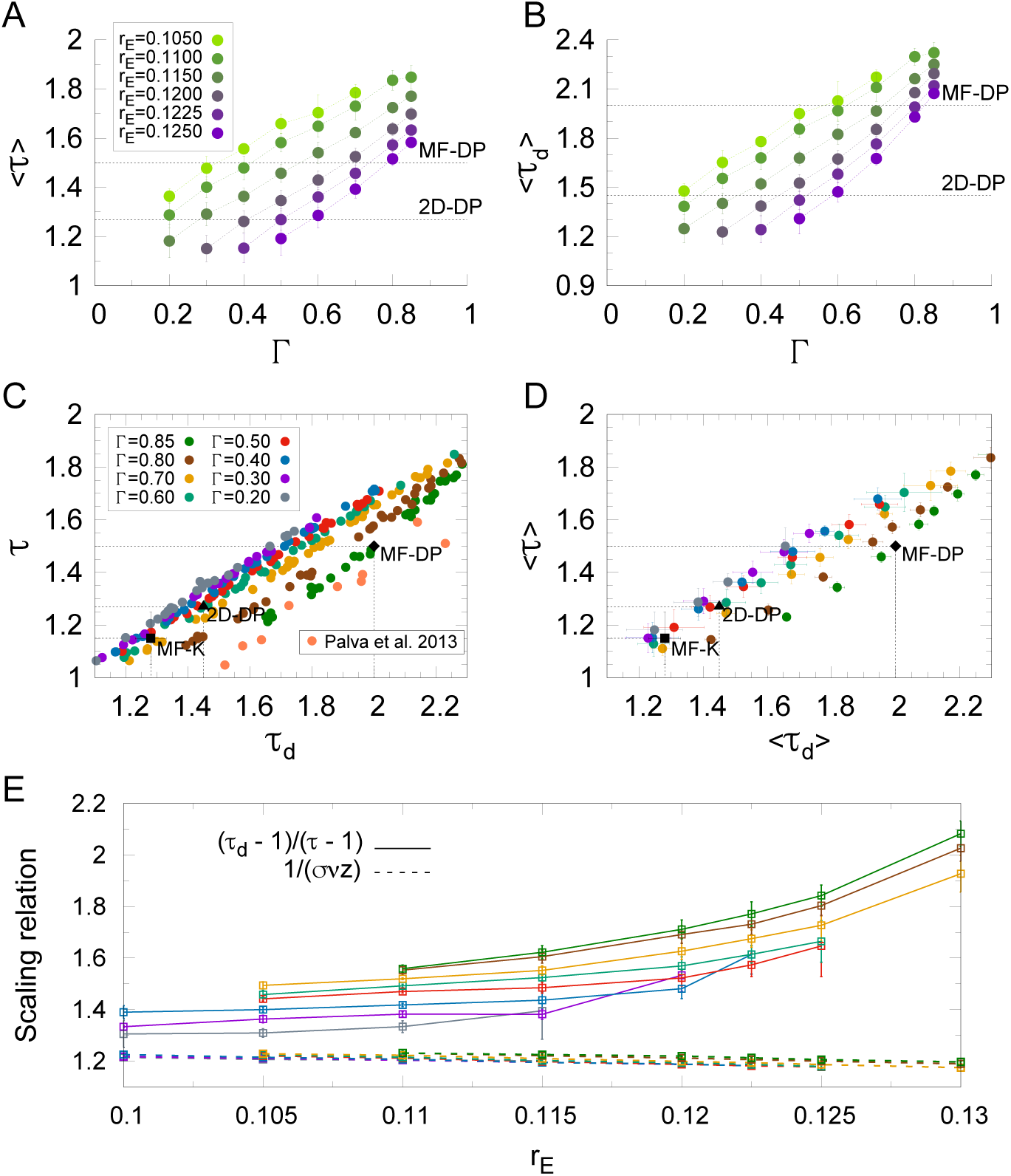
Dependence of the scaling behavior on the threshold defining an avalanche. (A) and (B) : exponents of avalanche size and duration, respectively, as a function of threshold parameter Γ (see Eq. 4). The same color code applies to (A) and (B). Exponents for avalanche size and avalanche duration (C) for 5 different runs and (D) averaged over the runs (the black points are the same as described in Fig. 6). (E) Left-hand (dashed line) and right-hand (solid line) sides of the scaling relation given by Eq. 7 as a function of *r*_E_ in the critical region. The color code in (C), (D) and (E) represents different values of Γ, as described in (C). In all figures, error bars represent the standard deviation over 5 runs.

Second, as the values of *τ* and *τ_d_* change with Γ, they keep a reasonably linear relationship with each other (Figs. 7C and 7D), which is consistent with the right-hand side of Eq. 7 being a constant. Moreover, and most importantly, this linear relationship depends on Γ. For increasing Γ, the linear spread of exponents in the (*τ*, *τ_d_*) plane is gradually displaced, eventually departing from the 2D-DP values and approaching the ones for MF-DP. Therefore, while for Γ ⋍ 0.5 it is possible to simultaneously find both 2D-DP exponents (*τ* ⋍ 1.268 and *τ_d_* ⋍ 1.45), for Γ ⋍ 0.85 it is possible to find both MF-DP exponents (*τ* = 1.5 and *τ_d_* = 2) at the critical region of the model.

Finally, changing the threshold parameter Γ does not fix the failure of the model to satisfy the scaling relation of Eq. 7 (Figs. 6F and 7E). While the measured exponent 1/(*σνz*) remains pretty stable across the critical region and regardless of Γ, the combination of avalanche exponents (right-hand side of Eq. 7) fail to reach the same value (Fig. 7E). In fact, as Γ increases (and exponents approach MF-DP values), the mismatch between the left- and right-hand sides of Eq. 7 worsens.

### Comparison with M/EEG experimental results

Regardless of the difficulties in reconciling a phase transition in the CROS model with a DP universality class, it is nonetheless interesting to point out that the model yields additional results that can be compared with experiments. For instance, the dependence of the average DFA exponent 〈*α*〉 on the exponents governing avalanche size and duration shows an inverted-U profile (Figs 8A and 8B, respectively), with a significant portion of the functions 〈*α*〉(〈*τ*〉) and 〈*α*〉(〈*τ_d_*〉) showing a decreasing behavior.

**Figure 8:**
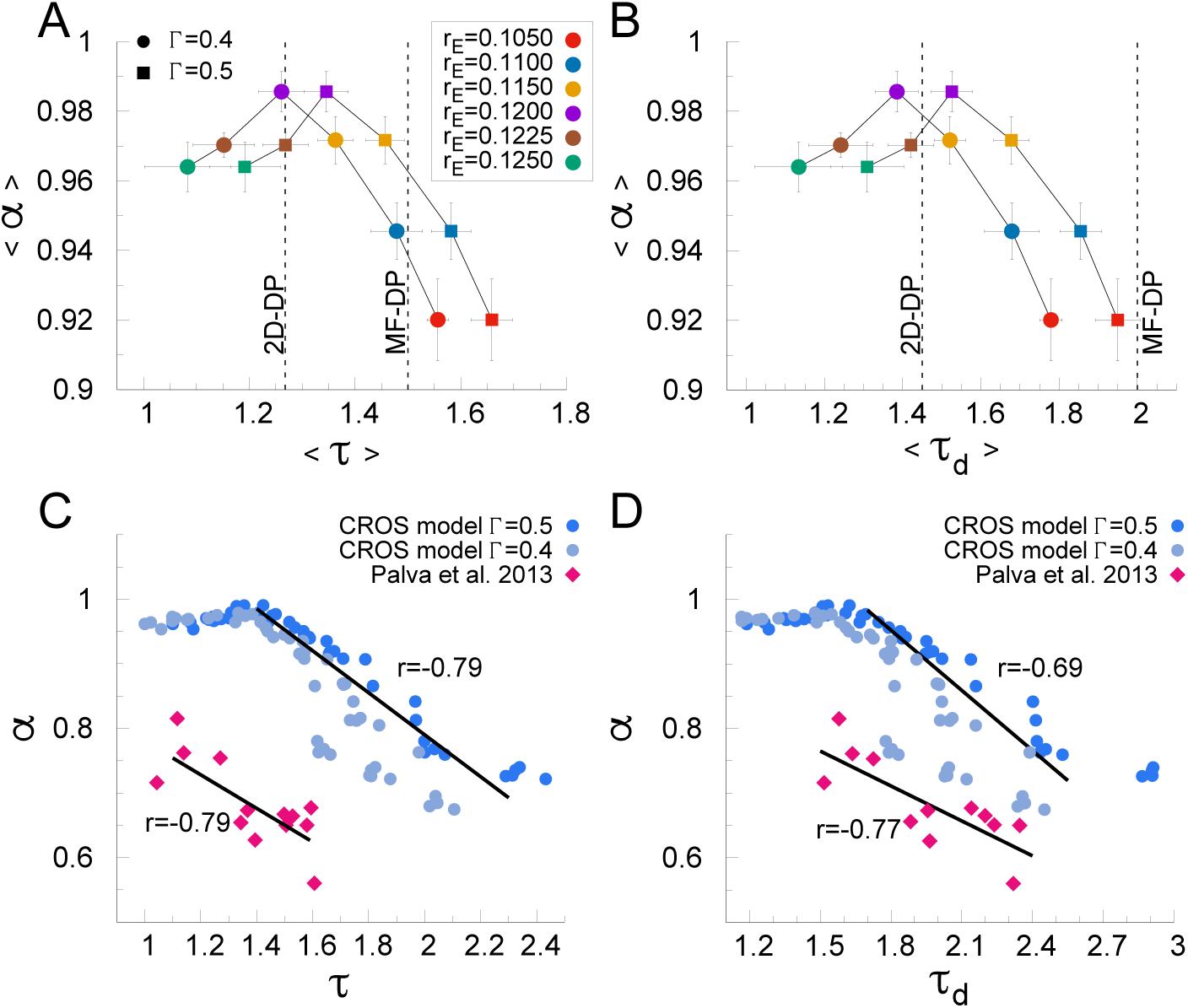
Similar between fingerprints of criticality in the CROS model and in M/EEG data. Relation between DFA exponent *α* and exponents for avalanche size (A) and duration (B), for different values of Γ, in the CROS model (in both cases, symbols and error bars respectively represent average and standard deviation over 5 realizations). Dashed lines show the theoretical exponents expected for 2D-DP and MF-DP universality classes. The same color code and dot type applies to (A) and (B). The negative correlation between *α* and *τ* (C) and *τ_d_* (D) on the CROS model (computed for Γ = 0.5) mimics the same tendency observed in M/EEG recordings of humans performing a threshold-stimulus detection task (data extracted from [20]). CROS model data showed in (C) and (D) holds for *r_E_* in the range [0.08 : 0.1250], whereas *r_I_* =0.60 was kept fixed.

This decreasing behavior is similar to what is observed experimentally in magneto- and electroencephalographic (M/EEG) recordings of humans performing a threshold-stimulus detection task [20]. Palva *et al* have found that source-reconstructed M/EEG data exhibit robust power-law long-range time correlations (i.e. well characterized DFA exponents) *as well as* scale-invariant avalanches (obtained by converting the recordings into a binary point process via a threshold at three standard deviations [20]).

It is important to emphasize the difference in time scales: while avalanches can last at most a couple of hundred milliseconds, long-range time correlations in the M/EEG data were observed for tens of minutes [20]. Yet, exponents characterizing the dynamics at such different scales were not independent, but rather negatively correlated, as shown in Figs 8C and 8D. Similar results are observed for recordings of subjects at rest, as well as for the dependence of the DFA exponent of behavioral time series (hits and misses) [20].

This very interesting experimental result, which connects signatures of criticality at very different time scales, can be partially reproduced by the CROS model. If one looks at single runs instead of averages (assuming that the result for a given experimental subject would be modeled by a single realization of the model disorder) and relaxes the loglikelihood ratio test requirement (therefore allowing “dirty power laws”), simulations within an interval of parameter values around the critical region give rise to DFA and avalanche exponents negatively correlated (Figs 8C and 8D). Moreover, the values of the correlations are very close to those observed experimentally. The main discrepancy between model and experiments is observed in the values of the DFA exponents, which are larger in the model. The quantitative agreement between the model and the M/EEG data can be marginally improved by decreasing the threshold parameter Γ (Figs. 8C and 8D).

Finally, it is interesting to note that the M/EEG data spreads linearly on the (*τ*, *τ_d_*) plane (Fig. 7E), a fact originally unreported [20]. Again, this is consistent with a constant right-hand side of Eq. 7. It remains to be verified whether the scaling between avalanche size and duration (Eq. 6) also holds, and whether the two sides of Eq. 7 match. In this context where one disregards the DFA exponents, the CROS model is able to *quantitatively* reproduce the spread of the avalanche exponents in the (*τ*, *τ_d_*) plane. As shown in Fig. 7E, the agreement between the model and the M/EEG data is best for larger values of the threshold parameter Γ.

## Conclusions

Given that the connection between critical phenomena and scale-invariant statistics has long been established, power-law distributions of avalanches should be a reasonable signature to look for if one believes a neuronal system is critical. It is important to keep in mind, however, that critical exponents depend on many factors, such as dimensionality, disorder and the nature of the phase transition. In particular, the avalanche size exponent *τ* ⋍ 1.5 originally observed by Beggs and Plenz [2] coincides with any model in the MF-DP universality class. But other experimental setups have revealed different exponents [9, 30, 35]. Likewise, models with different topologies or ingredients in their dynamics might give rise to exponents which differ from those of MF-DP.

Models with disorder and without an analytical solution, such as the one we studied here, can be particularly challenging. The CROS model is two-dimensional, with disorder, and the onset of its oscillations in principle seems physically different from the absorbing-active phase transition of the DP universality class. However, if one insists on the mean-field DP values and looks for the *τ* ⋍ 1.5 and *τ_d_* ⋍ 2 exponents, one may find one or the other, as we did by using the *κ* indexes as the main markers of criticality (Figs 4B and 5B). These, however, proved to be insufficient signatures. For instance, fixing the threshold parameter at Γ = 0.5, the MF-DP exponents could not be obtained simultaneously (Figs 6D and 6E). What we did obtain simultaneously for that threshold value were the size and duration exponents of 2D-DP (Figs 6D and 6E), a result that might seem reassuring given that the model is indeed two-dimensional and assuming that a connection between directed percolation and the onset of oscillations can be established at all. But even that shred of consistency disappears if the threshold is increased: for Γ ⋍ 0.85 we did obtain MF-DP exponents simultaneuously. In fact, as shown in Fig. 7C, it is possible to cover a significant fraction of the (*τ*, *τ_d_*) plane by varying two parameters, so it would be no surprise if the model could reproduce 1D-DP or 3D-DP as well. Recently, di Santo *et al* have investigated a different model with a similar claim, namely, that power-law distributed avalanches occur at the edge of synchronization [8]. They also simulated a twodimensional model, yet exhibited MF-DP avalanche exponents. It would be interesting to check what the strategies presented here would reveal if applied to their model.

Although the two-dimensional CROS model can exhibit exponents for avalanche size (*τ*) and duration (*τ_d_*) which are compatible with DP, we did not obtain a fully satisfactory connection between the two scenarios. In a DP-like phase transition, *τ* and *τ_d_* are expected to satisfy a scaling relation that involves the dependence of the average avalanche size 〈*s*〉 on the duration *d* (equations 6 and 7). This scaling relation is often satisfied experimentally [9, 30] and, although we found Eq. 6 to be valid within the critical region of the CROS model, we did not find a region in parameter space where Eq. 7 held. A number of details of the model could be further explored to investigate whether this issue can be circumvented, ranging from the very definition of an avalanche to a more sophisticated detection of the phase transition.

In any case, a theoretical understanding of whether DP can be an “effective theory” for the phase transition leading to oscillations is still missing. Consider, for instance, a recent study by Coleman *et al* that has numerically investigated avalanches in a variant of the globally coupled version of the paradigmatic Kuramoto model [6]. Tuning the large-*N* model to the slightly subcritical regime, the authors define avalanches as excursions of the Kuramoto order parameter between consecutive zero crossings. Through scaling analysis, they arrive at avalanche exponents *τ* = 1.15 ± 0.1 and *τ_d_* = 1.28 ± 0.05 [6]. Note that, on the one hand, these values are far from those of the MF-DP universality class, despite the mean-field nature of the model. Interestingly, on the other hand, their exponents are compatible with those we observed in the critical region of the two-dimensional CROS model for Γ = 0.5 (Figs 6D and 6E). In light of these results, therefore, it seems theoretically unwarranted to expect the “classical” MF-DP exponents (*τ* = 3/2 and *τ_d_* = 2) at the onset of collective oscillations.

Finally, we have shown that the CROS model is a very promising starting point for modeling some properties of M/EEG data. The model quantitatively reproduces the linear relation between the avalanche exponents (Fig. 7C), while their correlation with the DFA exponent is qualitatively reproduced (Fig. 8C and 8D). Given that the model overestimates long-range temporal correlations (as compared to the data), further investigation could delve into the question of which ingredients need to be added or modified in the model with the goal of attaining quantitative agreement. Furthermore, since the agreement with the data occurs near the phase transition of the model, it would be worth searching for dynamical mechanisms which could lead the system to self-organize around the critical region. These questions are beyond the scope of this manuscript and will be pursued elsewhere.

## Acknowledgments

We thank Renê Montenegro Filho, Ronald Dickman, Tiago L. Ribeiro, Ludmila Brochini and Pedro Carelli for useful discussions and comments. Financial support by Brazilian agencies FACEPE (grant IBPG-0335-1.05/14), CAPES (grant PVE 88881.068077/2014-01), CNPq (grant 310712/2014-9) and FAPESP Center for Neuromathematics (grant 2013/07699-0, São Paulo Research Foundation) is gratefully acknowledged.

